# Deep Learning-based Pseudo-Mass Spectrometry Imaging Analysis for Precision Medicine

**DOI:** 10.1101/2022.04.29.490098

**Authors:** Xiaotao Shen, Wei Shao, Chuchu Wang, Liang Liang, Songjie Chen, Sai Zhang, Mirabela Rusu, Michael P. Snyder

## Abstract

Liquid chromatography-mass spectrometry (LC-MS) based untargeted metabolomics provides systematic profiling of metabolic. Yet its applications in precision medicine (disease diagnosis) have been limited by several challenges, including metabolite identification, information loss, and low reproducibility. Here, we present the deepPseudoMSI project (https://www.deeppseudomsi.org/), which converts LC-MS raw data to pseudo-MS images and then processes them by deep learning for precision medicine, such as disease diagnosis. Extensive tests based on real data demonstrated the superiority of deepPseudoMSI over traditional approaches and the capacity of our method to achieve an accurate individualized diagnosis. Our framework lays the foundation for future metabolic-based precision medicine.

---

Liquid chromatography-mass spectrometry (LC-MS)-based untargeted metabolomics is a powerful tool that enables the identification of biomarkers for precision medicine^1^, such as diagnosing diseases^2^, customizing drug treatments^3^, and monitoring therapeutic outcomes^4^. The traditional processing and analysis method for LC-MS-based untargeted metabolomics in precision medicine can usually be divided into four steps^5^ (**Fig. S1**): (1) raw data processing, (2) data cleaning, (3) metabolite identification, and (4) diagnosis (prediction) model building. However, existing approaches suffer from several limitations. The first disadvantage is the information loss and misidentification of metabolites. Metabolite annotation is still one of the most challenging tasks for LC-MS-based untargeted metabolomics^5^. Most of the metabolite identification methods are based on the database resources, therefore, many metabolites not identified before are usually bypassed by the studies^6^. Current instruments usually detect tens or hundreds of thousands of metabolic features, however, only about 10% of those detected features could be identified in most experiments^6^. In addition, peak picking may lose low-intensity signals or mistakenly align features. This means that most of the information is lost in the further step of diagnosis/prediction model construction. The second disadvantage is the low reproducibility of LC-MS analysis^7^. During data acquisition, the retention time (RT), the mass-to-charge ratio (*m/z*), and signal intensity drift can commonly cause unwanted variations and significantly affect the diagnosis (prediction) accuracy. These substantially limit the application of LC-MS-based untargeted metabolomics in precision medicine^8^.

To overcome these limitations of the prior traditional methods, we presented the deepPseduoMSI project (**deep**-learning-based **Pseudo**-**M**ass **S**pectrometry **I**maging, https://www.deeppseudomsi.org/). Mass Spectrometry Imaging (MSI) can image thousands of molecules in a single experiment, making it a valuable tool for diagnosis^9^. The LC-MS raw data can be seen as an image containing millions of data points defined by retention time, mass-to-charge ratio, and intensity. Instead of peak picking to extract the metabolic feature table, we could also process the raw data as images to be handled by deep learning methods^10^.

The deepPseudoMSI includes two parts. The first part is the pseudo-MS image converter, which converts the LC-MS raw data to images (**Fig. 1a** and **Fig. S2**). The LC-MS raw data usually contains millions of data points, so we need to divide it into different pixels (or grids) based on the revolution in the x-axis (retention time) and y-axis (mass-to-charge ratio) to reduce the size. Briefly, all the data points in the same pixel are combined to represent the intensity of this pixel. Then, the intensity of each pixel is linearly transformed to the color of the pixel. Finally, one LC-MS raw data with millions of data points is converted into an image with thousands of pixels based on the resolution (for example, 224 × 224). The final generated “image” contains all the information from the LC-MS raw data, which is termed the pseudo-MS image. The second part is the pseudo-MS image predictor, a pre-trained VGG16 network (convolutional neural networks)^11^, which is fine-tuned to extract various image features from the pseudo-MS images to construct a prediction model (**Fig. 2b and Fig. S3**). Supervised deep learning models require a large number of labeled data to train^12^. To enlarge the number of pseudo-MS images for training, we adopt a strategy called data augmentation^13^ (**Fig. S4**). Briefly, we randomly add the RT, *m/*z, and intensity errors for each pseudo-MS image to simulate the drift during the data acquisition. Finally, several simulative images could be generated from one actual pseudo-MS image, which can significantly enlarge the number of images for training.

**Fig. 1.**
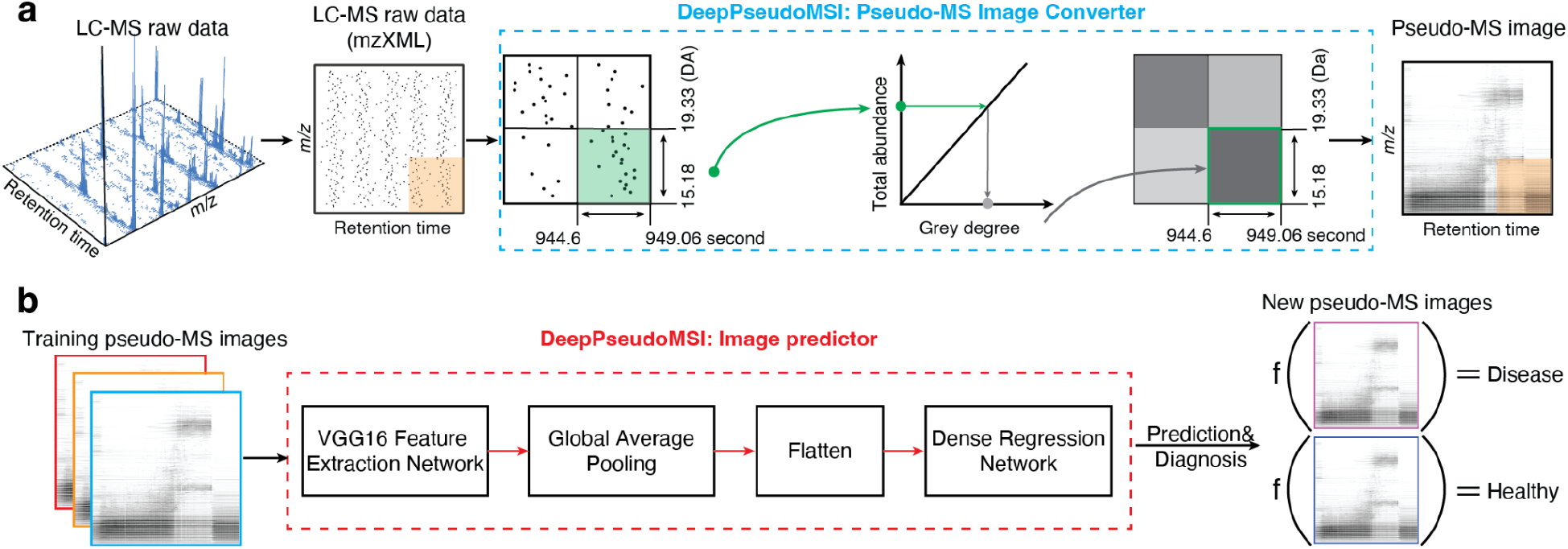
The workflow of converting LC-MS raw data to pseudo-MS images and the deep learning-based prediction model (deepPseudoMSI). **a**, Schematic of converting LC-MS raw data to pseudo-MS images (Image converter). LC-MS untargeted metabolomics raw data with millions of data points (x-axis represents RT, and the y-axis represents *m/z*) is binned into different pixels according to revolutions. The total intensity is calculated and transferred to a responded grey degree for each pixel. **b**, Schematic of prediction model construction (Image predictor). To generate more Pseudo-MS images for training, RT, *m/z*, and intensity drift are utilized for data augmentation for each pseudo-MS image. Then, the pseudo-MS images are projected for model training and construction using the VGG16 network.

**Fig. 2.**
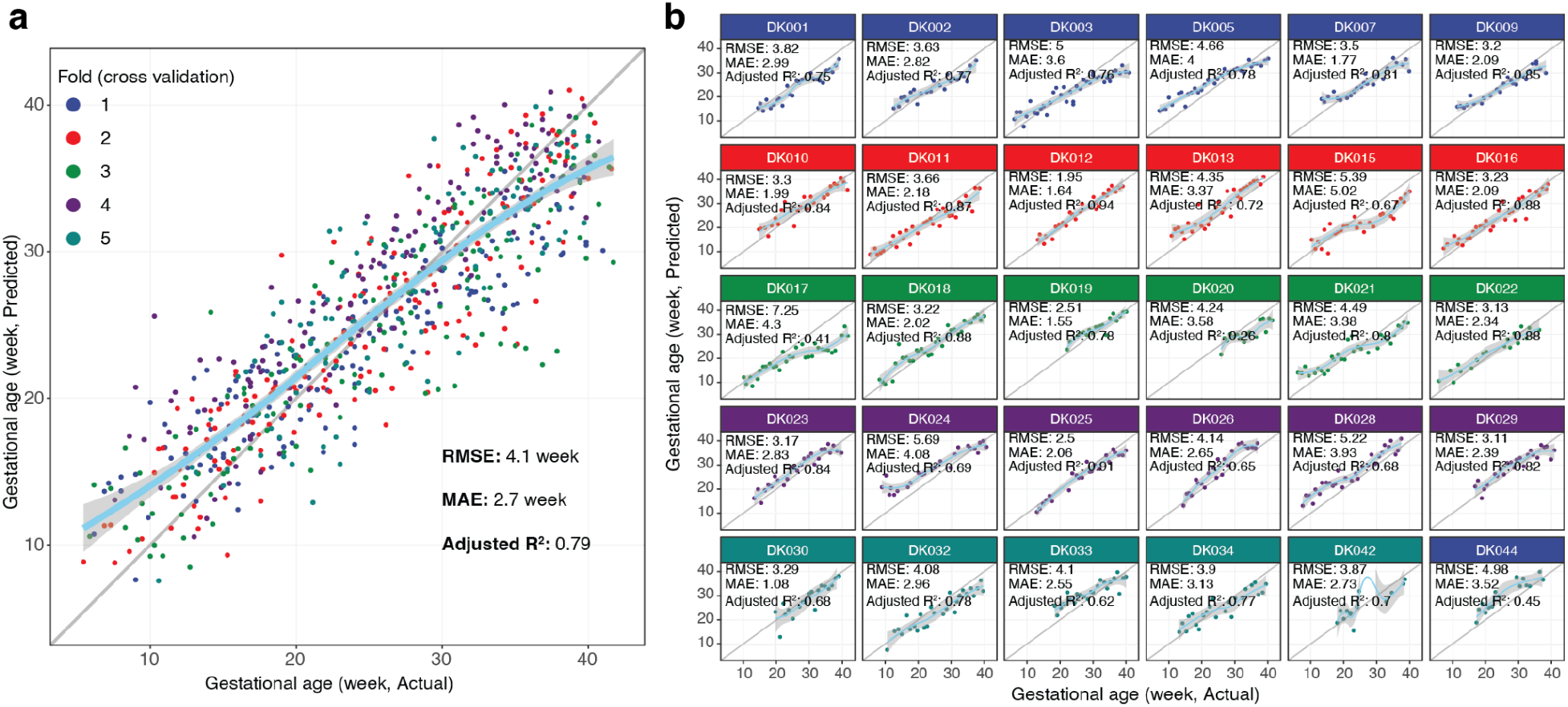
DeepPseudoMSI predicts gestational age in pregnant women. **a**, Gestational age predicted by deepPseudoMSI (y-axis) highly correlates with clinical values determined by the standard of care (x-axis). Different colors represent samples in different folds (5-fold cross-validation). **b**, Highly correlated GA predicted by deepPseudoMSI (y-axis) and actual GA (x-axis) at the individual level.

Compared to the traditional method, deepPseudoMSI does not need to annotate metabolites because all the information from the raw data is used for subsequent processing and analysis. Additionally, the drift of RT and *m/z* during data acquisition represents the shift of one pseudo-MS image on the x and y-axis. And the drift of intensity just represents the brightness changing of one pseudo-MS image. Our results show that the deep learning model can easily handle those variations and does not affect its prediction accuracy. Collectively, the pseudo-MS image can overcome the disadvantages of the traditional method, which may improve the application of LC-MS in precision medicine.

To gauge the effectiveness of deepPseudoMSI, it is used to predict the gestational age (GA, week) of pregnant women^14^ (**Fig. S5**) using our previously published dataset. This provides a more cost-effective method for pregnancy dating. First, the LC-MS raw data were converted to pseudo-MS images using the Pseudo-MS image converter. To identify the optimal resolution of the pseudo-MS images, we compared the generally used 224×244 and 1024×1024 resolutions presetting. And the first one achieved a better prediction result (RMSE: 3.61 *vs*. 6.10) (**Fig. S6**), so the 224×224 resolution was chosen for the pseudo-MS image generation. The data augmentation method was utilized to get lots of simulative pseudo-MS images for training to construct the prediction model. And then, the prediction model was built using the pseudo-MS image predictor. To evaluate the prediction model’s performance based on deepPseudoMSI, the 5-fold cross-validation method was utilized (**Fig. S7**). Intriguingly, the root mean square error (RMSE) is 4.1 weeks (mean absolute error (MAE) is 2.7 weeks. Adjusted R^2^ is 0.79) (**Fig. 2a**), which is better than the prediction result using the traditional method with all features (Random Forest model, RMSE: 4.34 weeks; adjusted R^2^: 0.76. **Fig. S9**. The permutation test *p*-value < 0.05). In addition, the deepPseudoMSI can get good prediction accuracy at the individual level (**Fig. 2b** and **Fig. S8**). This result demonstrates that the deepPseudoMSI has the potential to be leveraged for clinical diagnosis in the future.

To demonstrate that deepPseudoMSI can overcome the disadvantages of the traditional methods for LC-MS data, we designed an experiment to simulate the pervasive issue in LC-MS data acquisition, RT drift. Briefly, the random RT error was added to each raw data to simulate the RT drift during data acquisition (**Fig. 3a** and **Fig. S10**). We named the raw dataset “original dataset”, and the simulative dataset “RT drift dataset”. And then, both datasets were used for the raw data processing (traditional method) and pseudo-MS image conversion (deepPseudoMSI), respectively. The overlapped features between the original and the RT drift datasets are tiny (Jaccard index: 0.324, **Fig. 2b**), which is within the expectation^15^. Then we used the traditional method and deepPseudoMSI to construct the prediction model and validate results in original and RT drift datasets, respectively. Remarkably, the deepPseudoMSI has no difference in the prediction accuracy between the original and RT drift datasets (**Fig. 3c** and **Fig. 3d**). However, for the traditional method, the RT drift dataset’s prediction accuracy significantly decreases compared to the original dataset (**Fig. 3d**). About 16% of samples whose prediction errors are between 0-2 weeks in the original dataset then increased to 2-5 weeks in the RT drift dataset. Collectively, those results demonstrate that the deepPseudoMSI can overcome the disadvantages of the traditional methods for LC-MS-based untargeted metabolomics in diagnosis.

**Fig. 3.**
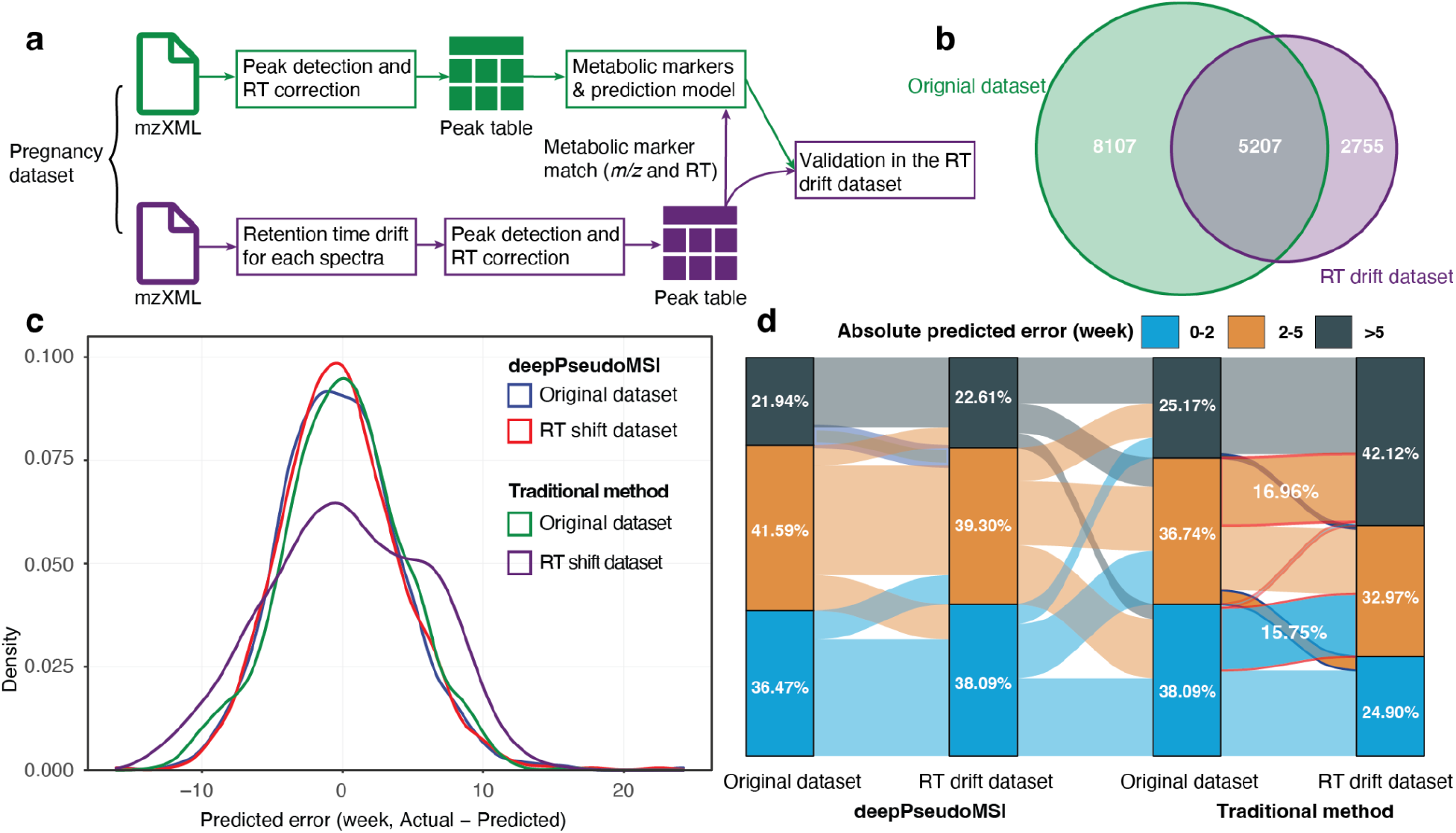
deepPseudoMSI can handle most of the disadvantages of the traditional method. **a**, Schematic simulation of RT drift in untargeted metabolomic data and then utilize the traditional method to process and construct prediction models. **b**, Venn diagram shows the metabolic features matching between original and RT drift datasets. **c**, Predicted error distribution of original and RT drift datasets that processed utilized deepPseudoMSI and traditional methods, respectively. **d**, Sankey diagram shows the absolute predicted errors for each sample in different datasets and methods.

To our best knowledge, this is the first systematic study that converts the LC-MS-based untargeted metabolomics data to pseudo-MS images and then takes advantage of the power of deep learning in image processing for precision medicine^16,17,18,19^. We also demonstrate that the deepPseudoMSI can overcome the limitations of the traditional method for LC-MS data in precision medicine. In summary, those results indicate that the deepPseudoMSI has the potential ability to significantly increase the application of mass spectrometry in clinics for precision medicine.

As a pilot study, our research has some shortcomings that we need to improve. First, deep learning methodology is a black-box-like process, and we don’t know the details of the pseudo-MS image process that contributes the most to our prediction. Second, we only use one mode of the LC-MS data (positive mode) to convert it to the pseudoMS image. Next, we plan to explore how to combine datasets of different chromatography and ESI modes to increase the prediction accuracy. We believe the deepPseudoMSI can provide a new data analysis direction for precision medicine using LC-MS-based untargeted metabolomics data. We only used untargeted metabolomics to demonstrate the application of deepPseudoMSI, this strategy can also be easily applied to LC-MS-based untargeted lipidomics and proteomics data.

## Methods

### PseudoMS image converter

The pseudo-MS image converter is designed and developed to convert the LC-MS-based untargeted metabolomics raw data to pseudo-MS images. Briefly, the LC-MS-based untargeted metabolomics raw data (from mass spectrometry instrument) is first converted to mzXML format data using msConvert^20^ or massconverter^21^. And then, the mzXML format data is imported to the R environment using the readMSData function from the MSnbase package^22^. Then the data points are filtered based on the *m/z*, RT, and intensity. The thresholds for the filtering should be based on the experiment and design. In our case study, the RT cutoff is set as RT > 50 and RT < 1000 seconds, and the *m/z* cutoff is set as *m/z* > 70 and *m/z* < 1000 Da. We then divide the data points by the y-axis (*m/z*) into different pixels (or grids) based on the set resolution. For example, if the pseudo-MS image resolution is set as 224 × 224, the data points in each scan are divided into 224 grids, and the data points in the same grid are combined as one pixel. The data points in one pixel have close retention time and mass-to-charge ratio, so they may be similar metabolites with the same biological functions. Then for the x-axis (RT), the scans are divided into different grids based on the resolution. Then the LC-MS raw data is converted into an image with thousands of pixels. For each pixel, it contains data points that are in the range of the pixel (x-axis and y-axis). Then the intensity of all the data points is log-transformed to correct heteroscedasticity and promote the low-intensity data point contribution^23^. The mean value of all the data points in this pixel is calculated to represent the pixel’s intensity. To transform the intensity of each pixel to color, we linearly transform the intensity of pixel to color (grey degree, from 0 - 255). Finally, the pseudo-MS image (black-and-white graph, png format) is generated with a specific resolution. The pseudo-MS image converter is written in R and available on GitHub (https://github.com/jaspershen/deepPseudoMSI/tree/main/code/pseudoMS-image-converter).

### Data augmentation for the training dataset

We developed an augmentation strategy to simulate pseudo-MS images for training. Briefly, for each mzXML format data, the MSnbase package is used to read it into the R environment. We randomly added an RT error, *m/z* error, and intensity error to all the data points in this spectrum. The RT error, *m/z* error, and intensity error are assigned, which are from the “error distributions”. For example, for the RT error, if we set it as 10 seconds, we will construct an “RT error distribution” (a normal distribution with a mean value of 10 seconds and an SD (standard variation) of 2 seconds). Then, for each data point in one scan, an RT error will be added randomly from the “RT error distribution”. The same strategy is used for *m/z* and intensity error adding. And then, the drifted mzXML data is converted to a pseudo-MS image using the pseudo-MS image converter. In the case study, we randomly generated 6 drifted pseudo-MS images for each data point.

### Pseudo-MS image predictor

The image predictor of deepPseudoMSI is a deep learning-based approach for predicting (diagnosis) using pseudo-MS images. Using the case study as an example, we first fine-tuned a pre-trained VGG16 network^11^ to extract various image features from the pseudo-MS images. The extracted image features were then fed into a global average pooling (GAP) layer, which transforms the input dimension from *N* × *N* × *C* to 1 × 1 × *C*, where *N* is the size of each feature image and *C* is the number of features. The output of the GAP layer was flattened and connected to a stack of three dense layers to regress the gestational age. One advantage of using the GAP layer is that it converts feature images of any dimension to 1×1, allowing our image predictor network to predict the gestational age from pseudo-MS images of any size. The GAP layer can also prevent the deep neural network from overfitting since it has significantly reduced the number of model parameters. We trained our neural network using 5,250 pseudo-MS images (including the drifted pseudo-MS images using a data augmentation strategy) from 30 subjects (750 samples) with a 5-fold cross-validation on the NVIDIA GeForce RTX 2080 GPU (8GB memory, 14,000 MHz clock speed). In training, we used the Adam optimizer with an initial learning rate of 0.0001 and a learning rate decay of 0.98. The batch size was set to be 8. The training was terminated after 100 epochs. The pseudo-MS predictor is written in Python and available on GitHub (https://github.com/jaspershen/deepPseudoMSI/tree/main/code/pseudoMS-image-predictor).

### Retention time (RT) drift dataset generation

All the mzXML format data were loaded using the MSnbase R package^22^. Then for each spectrum, the retention time (RT) was randomly added with a specific error to simulate RT drift in LC-MS data acquisition (RT error is 60 seconds and SD is 10 seconds, see the “Data augmentation for the training dataset” section). Then the RT drift data were subjected to peak detection and alignment using XCMS^24^, and the parameter setting is the same as in the “Data augmentation for the training dataset” section.

### Alignment of two metabolic peak tables

Two metabolic feature tables were aligned according to *m/z* and RT using the masstools package (mz_rt_match function) from the tidyMass project^21^. Briefly, only the features in two metabolic feature tables within the setting cutoff for *m/z* matching (< 10 ppm) and RT matching (< 30 seconds) are considered the same features. If one feature matches multiple features, only the feature with the minimum RT matching error remains.

### General statistics analysis and data visualization

All the general statistical analysis and data visualization are performed utilizing Rstudio (Version 1.3.959) and R environment (Version 4.1.2). Most of the R packages and their dependencies used in this study are maintained in CRAN (https://cran.r-project.org/), Bioconductor (https://www.bioconductor.org/), or GitHub. The detailed information on R packages is provided in the **Supplementary Note**. The R package ggplot2 (version 3.2.21) was used to perform all the data visualization in this study.

### Five-fold cross-validation

To avoid information leakage, all the 30 subjects are randomly assigned to 5 groups (*sample* function in R), and each group has six subjects. Then all the samples are assigned to different groups based on the subjects. So for each subject, all its samples are in the same group.

### Random Forest prediction model

The boruta algorithm^25^ (R package Boruta, version 6.0.0) is utilized to select potential biomarkers. Briefly, it duplicates the dataset and shuffles the values in each column. These values are called shadow features. Then, it trains a Random Forest classifier (R package randomForest) on the dataset and checks for each of the real features if they have higher importance. If it does, the algorithm will record the feature as “important”. This process is repeated 100 iterations. In essence, the algorithm is trying to validate the importance of the feature by comparing it with randomly shuffled copies, which increases the robustness. This is performed by comparing the number of times a feature did better with the shadow features using a binomial distribution. Finally, the confirmed features are selected as potential biomarkers for Random Forest model construction.

In the Random Forest model, all the parameters are used as default settings except ntree (number of trees to grow) and mtry (number of variables randomly sampled as candidates at each split). Those two parameters are optimized on the training dataset, they are combined to form a set. The performance of each set of parameters is evaluated using the mean squared error (MSE). The parameter pair with the smallest MSE is used to build the final prediction model.

We utilize the 5-fold cross-validation method to evaluate the prediction accuracy of our models. Briefly, it is selected as the validation dataset for each fold, and the remaining four-fold data are used for the training dataset. The training dataset is utilized to get the potential biomarkers using the feature selection method described above. Then a Random Forest prediction model is built based on the training dataset. Then the external validation model is utilized to demonstrate its prediction accuracy. The predicted GA and actual GA for the validation dataset are plotted to observe the prediction accuracy. Then the RMSE (root mean squared error), MAE (mean absolute error), and adjusted R^2^ are used to quantify the prediction accuracy.

For internal validation, the bootstrap sampling method is utilized^4^. We randomly sampled the same number of samples from the training dataset with replacement (about 63% of the unique samples on average). We then used it as an internal training dataset to build the Random Forest prediction model using the same selected features and optimized parameters. The remaining about 37% of the samples were used as the internal validation dataset. Those steps repeat 1,000 times. Finally, we got more than one predicted GA value for each sample. The mean value of multiple predicted GA values is used as the final average predicted GA and used to calculate RMSE, MAE, and adjusted R^2^.

### Permutation test

The first permutation test was utilized to calculate *p*-values to assess if the Random Forest prediction models are not overfitting. In brief, firstly, all the responses (GA, week in this study) are randomly shuffled for both training and validation datasets, respectively. Secondly, the potential biomarkers are selected, and the parameters of Random Forest are optimized in the training dataset using the method described above. Thirdly, the Random Forest prediction model uses the selected features and optimized parameters in the training dataset. Finally, we use this random forest prediction model to get the predicted responses for the validation dataset. Then we get the null RMSE and adjusted R^2^. We repeat this process 1,000 times, getting 1,000 null RMSE and 1,000 null adjusted R^2^ vectors. Using maximum likelihood estimation, these null RMSE values and adjusted R^2^ values are modeled as Gamma distribution, and then the cumulative distribution function (CDF) is calculated. Finally, the *p*-values for the real RMSE and adjusted R^2^ are calculated from the null distributions, respectively.

The second permutation test was utilized to calculate the *p*-value to assess if the depPseudoMSI performs better than the traditional method. In brief, for the traditional method, we randomly shuffled the subjects to different 5-folds and then used this to construct the Random Forest prediction model and get a new prediction result. This step was repeated 1,000 times, so we have 1,000 prediction results for the traditional model. Then the *p*-value was calculated based on the method described above.

### Sample preparation and data acquisition of case study

All the sample preparation and data acquisition for the case study can be found in our previous publication^14^. In brief, 30 pregnant women were recruited, and 750 blood samples were collected during the study. Then all the blood samples were processed for LC-MS analysis.

### LC-MS-based untargeted metabolomics raw data processing

The mzXML format data (RPLC positive mode) were placed into different folders according to their class (for example ‘‘Blank’’, ‘‘QC’’ and ‘‘Subject’’) and then subjected to peak detection and alignment using the massprocesser package from the tidyMass project^21^ based on XCMS^24^. Briefly, the peak detection and alignment were performed using the centWave algorithm^24^. The key parameters were set as follows: method = ‘‘centWave’’; ppm = 15; snthr = 10; peakwidth = c(5, 30); snthresh = 10, prefilter = c(3, 500); minifrac = 0.5; mzdiff = 0.01; binSize = 0.025 and bw = 5. Finally, the generated MS^1^ metabolic feature table (peak table) includes the mass-to-charge ratio (*m/z*), retention time (RT, second), peak abundances for all the samples, and other information. This MS^1^ metabolic feature table is used for the subsequent data cleaning using the masscleaner package from the tidyMass project^21^. Briefly, the features detected in less than 20% QC samples were removed as noisy from the metabolic feature table. Then the missing values (MV) were imputed using the k-nearest neighbors (KNN) algorithm. Then the metabolic feature table is used for subsequent statistical analysis.

## Data availability

The LC-MS data (mzXML format, RPLC positive mode) were deposited to the NIH Common Fund’s National Metabolomics Data Repository (NMDR) website, the Metabolomics Workbench, and the project ID is PR000918 (https://doi.org/10.21228/M81H58). The metabolic feature and pseudo-MS images are provided on the deepPseduoMSI project website (https://www.deeppseudomsi.org/#case_study), and the metabolic feature tables also are provided as **Supplementary Data 1** and **2**.

## Code availability

The code of deepPseudoMSI and all the code for data processing, statistical analysis, and data visualization in this study have been provided on GitHub (https://github.com/jaspershen/deepPseudoMSI) under the MIT license for noncommercial use. All the statistical analyses were written by R, also provided as **Supplementary Data 3**.

## Acknowledgments

We thank Dr. Axel Brunger for the advice on the manuscript.

## Author contributions

X.S. conceptualized the study. X.S. and M.P.S. conceived the method and supervised its implementation. X.S. developed the pseudo-MS image converter algorithm. W.S. and X.S. developed the pseudo-MS image predictor. S.Z. inspected the deep learning method. X.S. and C.W. built the websites for the project. X.S., L.L., and S.C. provided and prepared the case study data. X.S., W.S., and C.W. analyzed the case study data. X.S. and C.W. designed and made the figures. X.S, C.W., W.S., and M.S.P wrote the manuscript. All authors contributed to the reviewing and editing of the final manuscript.

## Competing interests

M.P.S. is a co-founder and member of the scientific advisory board of Personalis, Qbio, January, SensOmics, Protos, Mirvie, NiMo, Onza, and Oralome. He is also on the scientific advisory board of Danaher, Genapsys, and Jupiter. M.R. is a consultant for Roche. Other authors declare no conflict of interests.

## Additional information

**Correspondence and requests for materials** should be addressed to X.S. or M.P.S.

## Notes

https://www.deeppseudomsi.org/

## Reference

1. Wishart, D. S. Emerging applications of metabolomics in drug discovery and precision medicine. Nat. Rev. Drug Discov. 15, 473–484 (2016).

2. Masoodi, M. et al. Metabolomics and lipidomics in NAFLD: biomarkers and non-invasive diagnostic tests. Nat. Rev. Gastroenterol. Hepatol. 18, 835–856 (2021).

3. Kok, M., Maton, L., van der Peet, M., Hankemeier, T. & Coen van Hasselt, J. G. Unraveling antimicrobial resistance using metabolomics. Drug Discov. Today (2022) doi:10.1016/j.drudis.2022.03.015.

4. Jia, H. et al. Predicting the pathological response to neoadjuvant chemoradiation using untargeted metabolomics in locally advanced rectal cancer. Radiother. Oncol. 128, 548–556 (2018).

5. Alseekh, S. et al. Mass spectrometry-based metabolomics: a guide for annotation, quantification and best reporting practices. Nat. Methods 18, 747–756 (2021).

6. Chaleckis, R., Meister, I., Zhang, P. & Wheelock, C. E. Challenges, progress and promises of metabolite annotation for LC–MS-based metabolomics. Current Opinion in Biotechnology vol. 55 44–50 (2019).

7. Shen, X. et al. Normalization and integration of large-scale metabolomics data using support vector regression. Metabolomics vol. 12 (2016).

8. Cui, L., Lu, H. & Lee, Y. H. Challenges and emergent solutions for LC-MS/MS based untargeted metabolomics in diseases. Mass Spectrom. Rev. 37, 772–792 (2018).

9. Unsihuay, D., Mesa Sanchez, D. & Laskin, J. Quantitative Mass Spectrometry Imaging of Biological Systems. Annu. Rev. Phys. Chem. 72, 307–329 (2021).

10. Aggarwal, R. et al. Diagnostic accuracy of deep learning in medical imaging: a systematic review and meta-analysis. NPJ Digit Med 4, 65 (2021).

11. Simonyan, K. & Zisserman, A. Very Deep Convolutional Networks for Large-Scale Image Recognition. arXiv [cs.CV] (2014).

12. Sun, C., Shrivastava, A., Singh, S. & Gupta, A. Revisiting Unreasonable Effectiveness of Data in Deep Learning Era. 2017 IEEE International Conference on Computer Vision (ICCV) (2017) doi:10.1109/iccv.2017.97.

13. Khalifa, N. E., Loey, M. & Mirjalili, S. A comprehensive survey of recent trends in deep learning for digital images augmentation. Artif Intell Rev 1–27 (2021).

14. Liang, L. et al. Metabolic Dynamics and Prediction of Gestational Age and Time to Delivery in Pregnant Women. Cell 181, 1680–1692.e15 (2020).

15. Brunius, C., Shi, L. & Landberg, R. Large-scale untargeted LC-MS metabolomics data correction using between-batch feature alignment and cluster-based within-batch signal intensity drift correction. Metabolomics 12, 173 (2016).

16. Liu, T., Siegel, E. & Shen, D. Deep Learning and Medical Image Analysis for COVID-19 Diagnosis and Prediction. Annu. Rev. Biomed. Eng. (2022) doi:10.1146/annurev-bioeng-110220-012203.

17. Calivà, F. et al. Studying osteoarthritis with artificial intelligence applied to magnetic resonance imaging. Nat. Rev. Rheumatol. 18, 112–121 (2022).

18. Bera, K., Braman, N., Gupta, A., Velcheti, V. & Madabhushi, A. Predicting cancer outcomes with radiomics and artificial intelligence in radiology. Nat. Rev. Clin. Oncol. 19, 132–146 (2022).

19. Behrmann, J. et al. Deep learning for tumor classification in imaging mass spectrometry. Bioinformatics 34, 1215–1223 (2018).

20. Chambers, M. C. et al. A cross-platform toolkit for mass spectrometry and proteomics. Nat. Biotechnol. 30, 918–920 (2012).

21. Shen, X. et al. TidyMass: An Object-oriented Reproducible Analysis Framework for LC-MS Data. bioRxiv 2022.03.15.484499 (2022) doi:10.1101/2022.03.15.484499.

22. Gatto, L., Gibb, S. & Rainer, J. MSnbase, efficient and elegant R-based processing and visualisation of raw mass spectrometry data. doi:10.1101/2020.04.29.067868.

23. Blaise, B. J. et al. Statistical analysis in metabolic phenotyping. Nat. Protoc. 16, 4299–4326 (2021).

24. Smith, C. A., Want, E. J., O’Maille, G., Abagyan, R. & Siuzdak, G. XCMS: processing mass spectrometry data for metabolite profiling using nonlinear peak alignment, matching, and identification. Anal. Chem. 78, 779–787 (2006).

25. Kursa, M. B. & Rudnicki, W. R. Feature Selection with the Boruta Package. J. Stat. Softw. 36, 1–13 (2010).

