## Supplementary for "Deep Learning-based Pseudo-Mass Spectrometry Imaging Analysis for Precision Medicine"

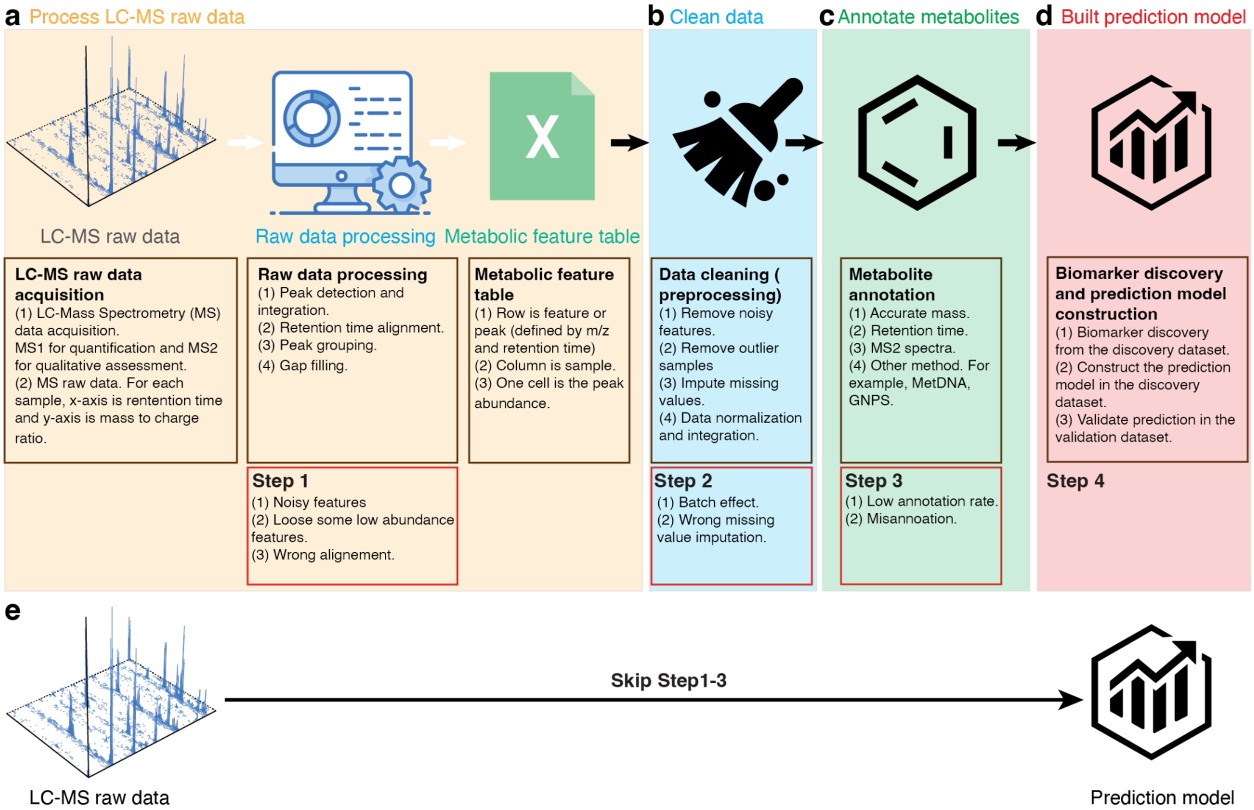


**Fig. S1. The traditional method for LC-MS-based untargeted metabolomics in diagnosis and precision medicine. a**, LC-MS raw data acquisition and raw data processing. Raw data processing contains peak picking, retention time correction and peak grouping. **b**, Data cleaning is used to remove the unwanted variations during sampling collection, sample preparation and data acquisition. **c**, Metabolite annotation is used to assign compound information for metabolic features from the metabolic feature table. The most commonly used method is to match the experimental features with metabolite standards. **d**, The last step is finding potential biomarkers and constructing the prediction model (diagnosis). **e**, To overcome the disadvantages of the traditional method, we could develop a new strategy that skips steps 1-3.

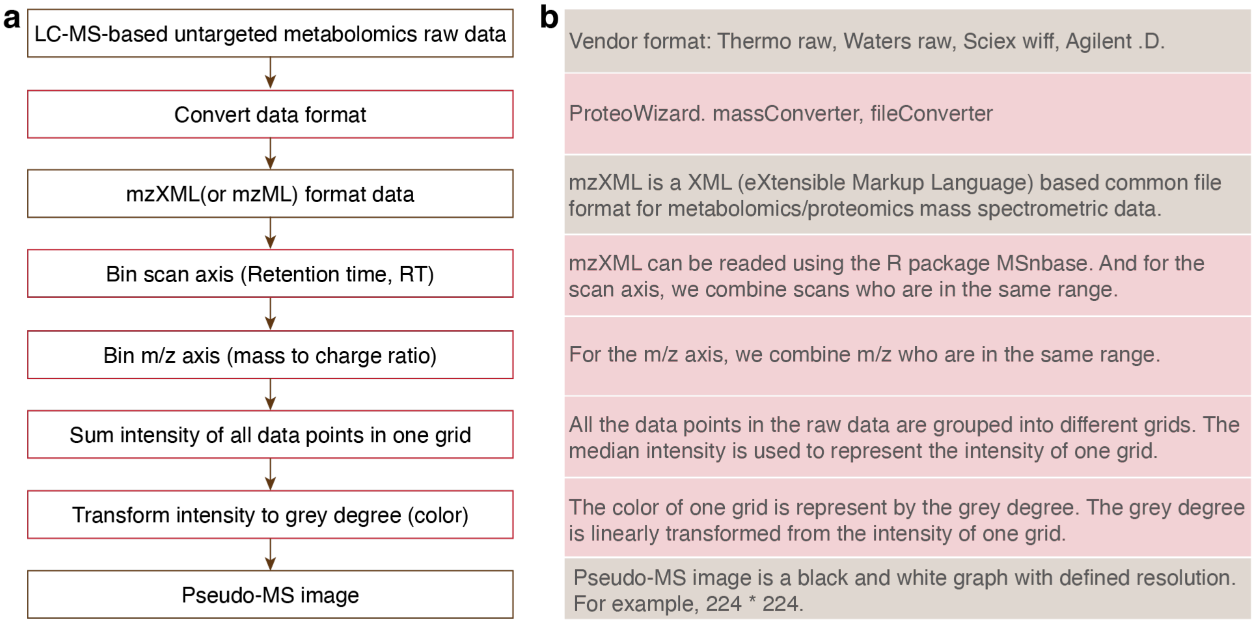


**Fig. S2. Workflow of pseudo-MS image converter. a**, The detailed steps for pseudo-MS image converter. **b**, The explanation for each step in **a**.

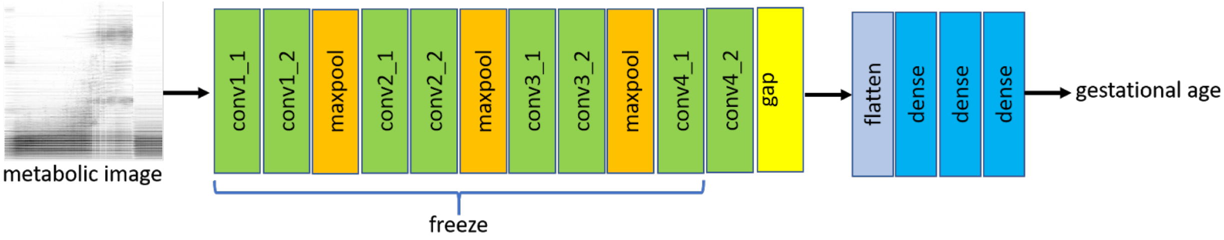


**Fig. S3. Workflow of pseudo-MS image predictor.** Deep neural network for gestational age prediction. The number of outputs of the three dense layers is 64, 8, and 1, respectively.

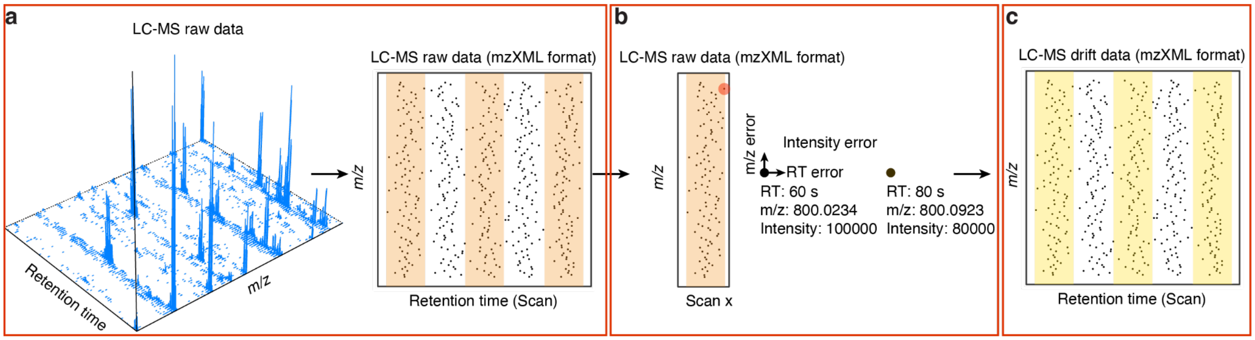

**Fig. S4. Workflow of pseudo-MS image augmentation. a**, LC-MS raw data is converted into mzXML format data. **b**, For each data point in LC-MS raw data, random RT error, *m/z* error, and intensity error are added to it. (**c**) Finally, one new pseudo-MS image is generated.

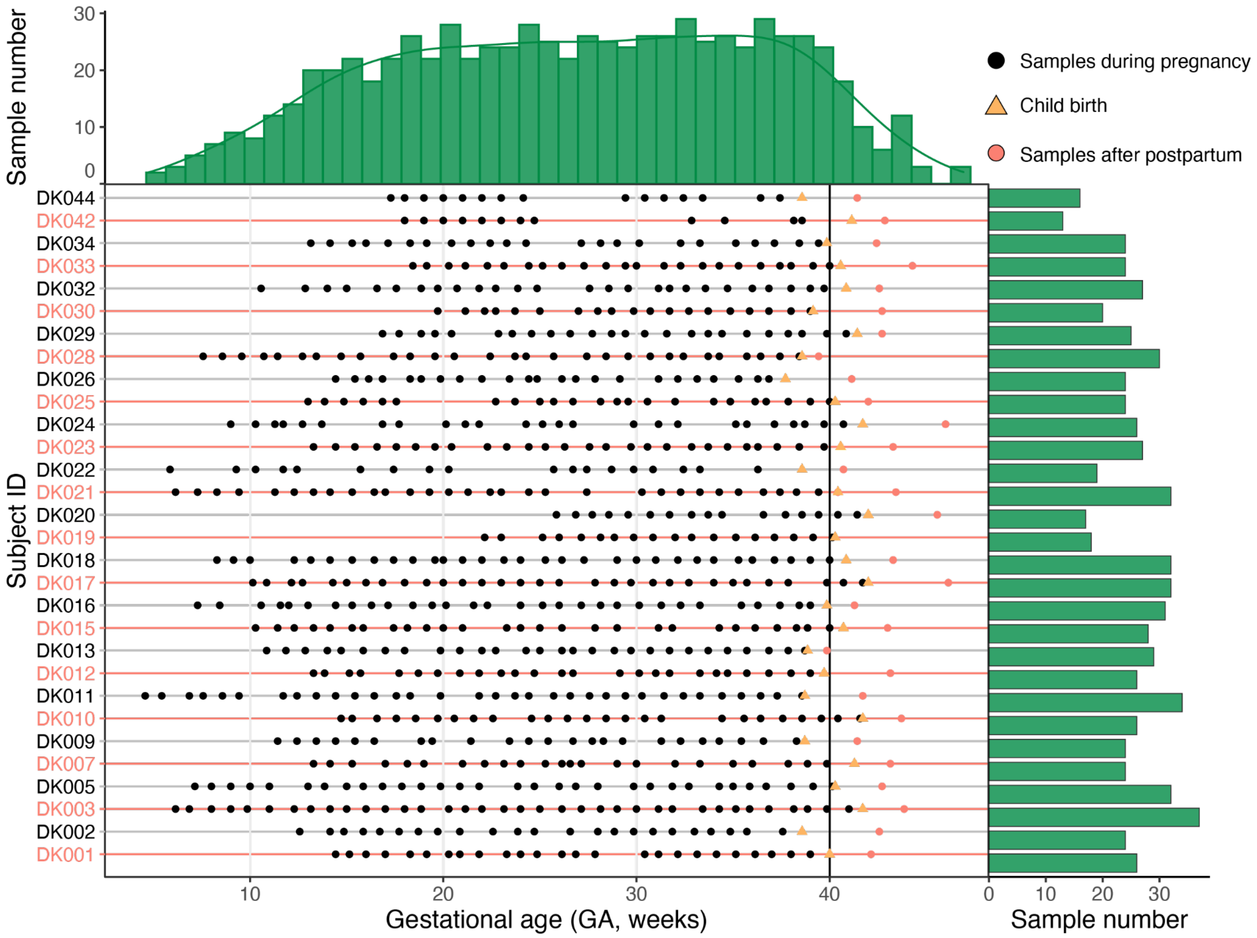


**Fig. S5. Case study overview.** The case study dataset is from our previous publication[1]. For each participant, we collected multiple blood samples during pregnancy.

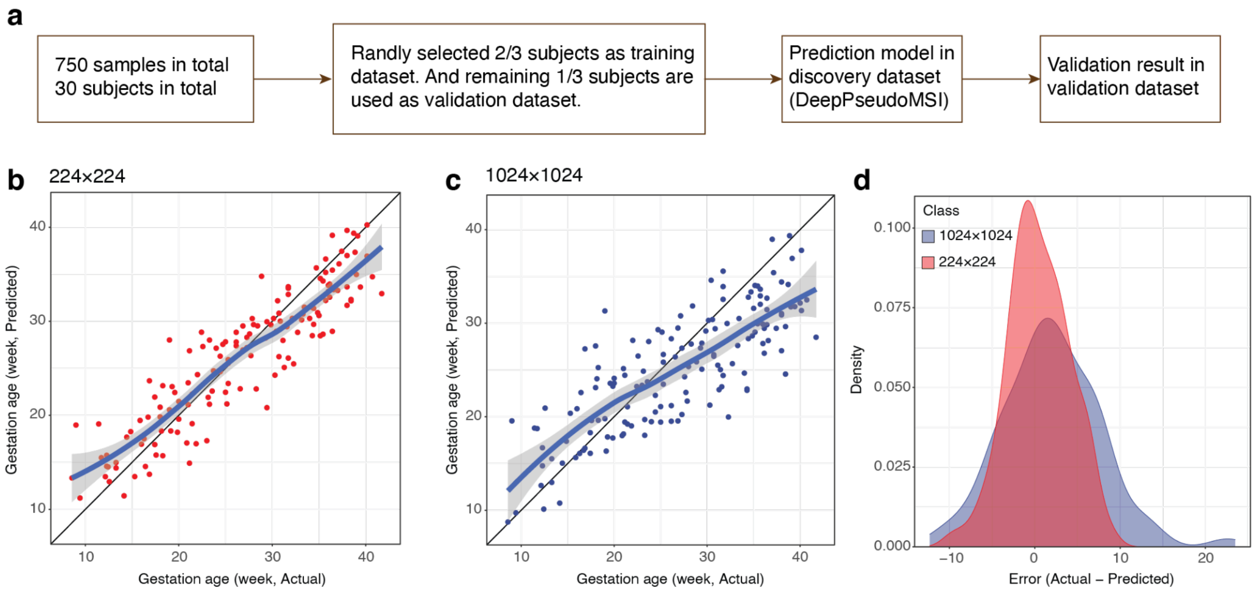


**Fig. S6. Pseudo-MS image resolution optimization for the case study. a**, The workflow shows how to use the case study to optimize the resolution of the pseudo-MS image. **b**, Actual GA (x-axis) versus predicted GA (y-axis) using the prediction model from resolution 224×224 (RMSE: 3.61, MAE: 2.44, Adjusted $R^2$: 0.83). **c**, Actual GA (x-axis) versus predicted GA (y-axis) using the prediction model from resolution 1024×1024 (RMSE: 6.10, MAE: 3.90, Adjusted $R^2$: 0.56). **d**, The prediction error distributions for prediction models from 224×224 and 1024×1024.

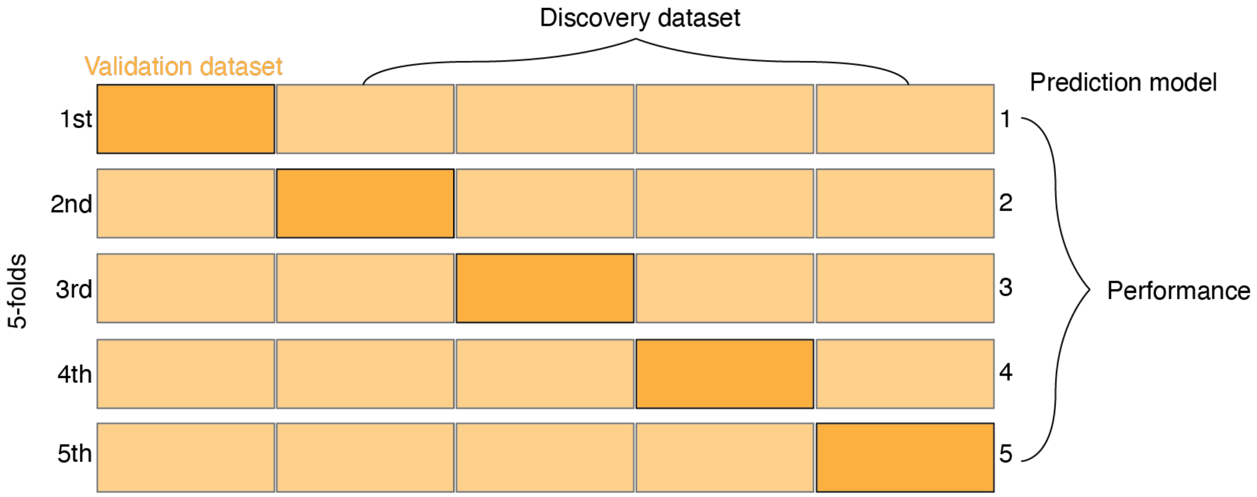


**Fig. S7. A five-fold cross-validation method was used in the case study.**

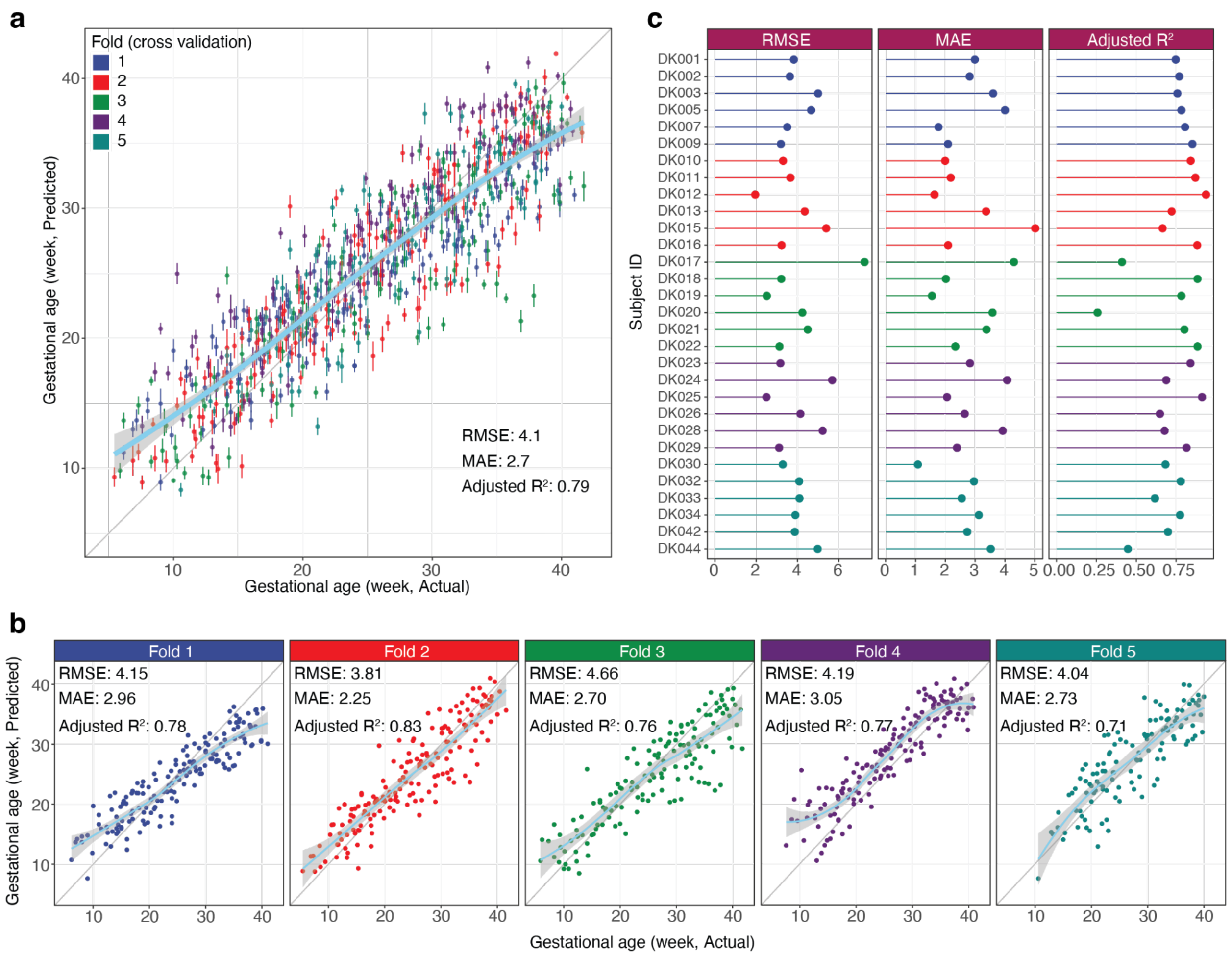


**Fig. S8. The prediction results in the case study using deepPseudoMSI. a**, Gestational age (GA, week) predicted by deepPseudoMSI (y-axis) highly correlates with clinical values determined by the standard of care (x-axis). Different colors represent samples in different folds (5-fold cross-validation). RMSE (root mean squared error), MAE (mean absolute error), and adjusted $R^2$ for each participant. For each sample, we also have internal validation (1,000 times) to give the range of prediction. **b**, The prediction results for each fold in the 5-fold cross-validation. **c**, The RMSE, MAE, and adjusted $R^2$ for each participant.

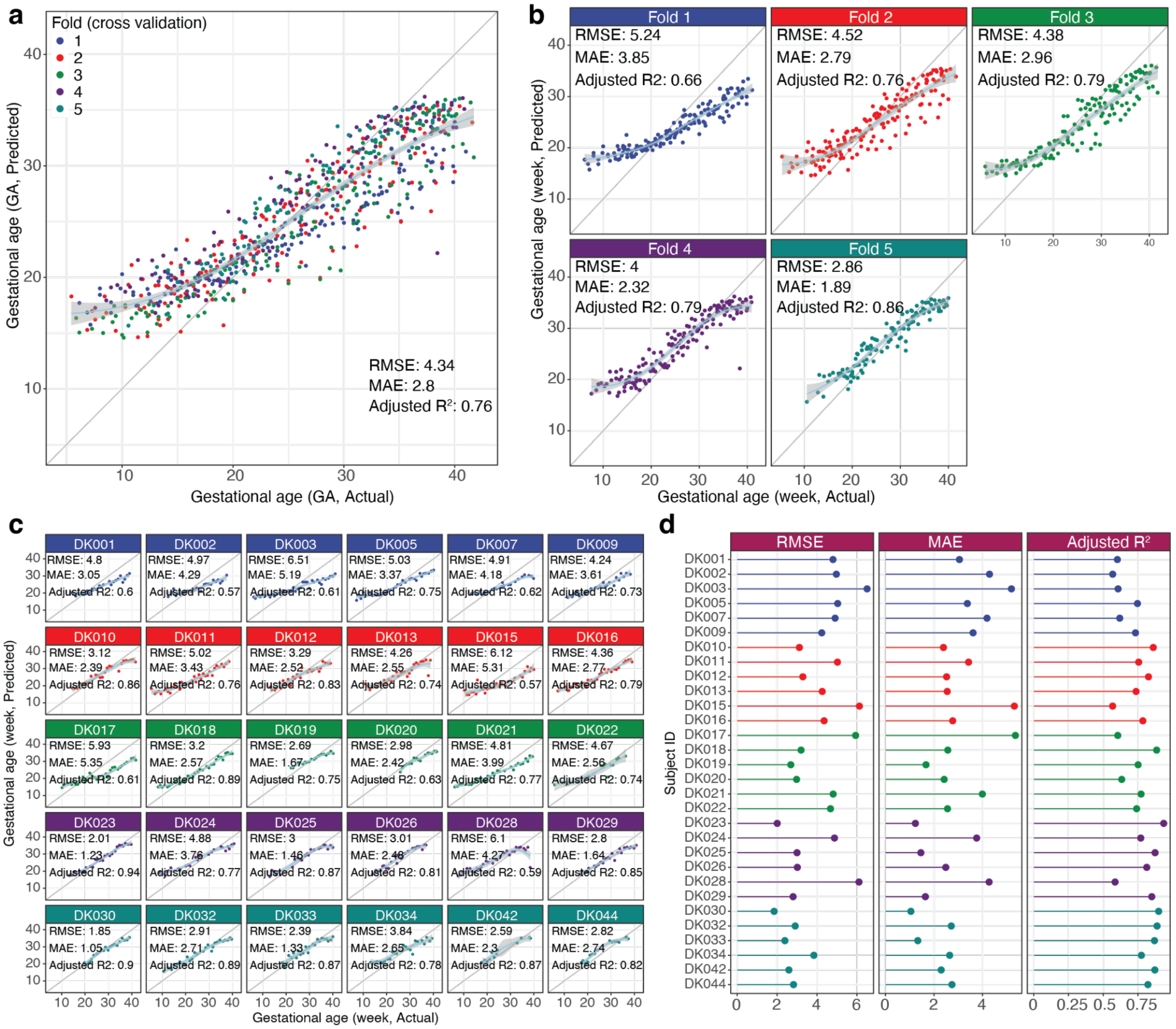


**Fig. S9. The prediction results in the case study using the traditional method. a**, Predicted gestational ages (GA, week) by the traditional method (y-axis) and the clinical values determined by the standard of care (x-axis). Different colors represent samples in different folds (5-fold cross-validation). **b**, The prediction results for each fold in the 5-fold cross-validation. **c**, Predicted GAs and actual GAs for each participant. **d**, The RMSE, MAE, and adjusted $R^2$ for each participant.

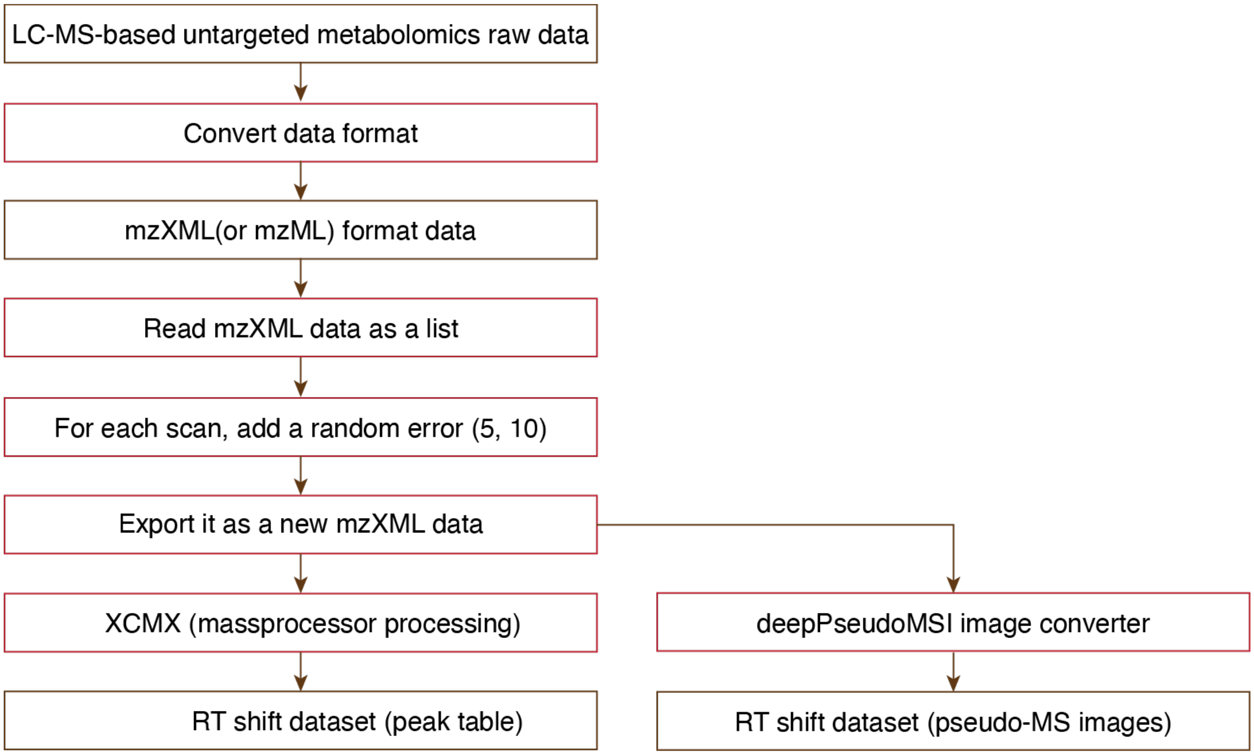

**Fig. S10. Simulation of RT shift in LC-MS-based untargeted metabolomics data acquisition.**

### Supplementary Data

**Supplementary Data 1.** Metabolic features table for the original dataset.
**Supplementary Data 2.** Metabolic feature table for the RT drift dataset.
**Supplementary Data 3.** Code for data processing, statistical analysis, and visualization.

### Supplementary Note

```
R version 4.1.2 (2021-11-01)
Platform: x86_64-apple-darwin17.0 (64-bit)
Running under: macOS Monterey 12.3

Matrix products: default
LAPACK: /Library/Frameworks/R.framework/Versions/4.1/Resources/lib/libRlapack.dylib

locale:
[1] en_US.UTF-8/en_US.UTF-8/en_US.UTF-8/C/en_US.UTF-8/en_US.UTF-8

attached base packages:
[1] grid      stats     graphics  grDevices utils     datasets  methods   base

other attached packages:
 [1] Boruta_7.0.0        randomForest_4.7-1  VennDiagram_1.7.1   futile.logger_1.4.3
 [5] magrittr_2.0.2      masstools_0.99.9    plyr_1.8.6          forcats_0.5.1.9000
```

```
 [9] stringr_1.4.0       dplyr_1.0.8         purrr_0.3.4         readr_2.1.2
[13] tidyr_1.2.0         tibble_3.1.6        ggplot2_3.3.5       tidyverse_1.3.1

loaded via a namespace (and not attached):
 [1] fs_1.5.2              ProtGenerics_1.26.0   lubridate_1.8.0       doParallel_1.0.17
 [5] httr_1.4.2            ggsci_2.9             MSnbase_2.20.4        tools_4.1.2
 [9] backports_1.4.1       utf8_1.2.2            R6_2.5.1              affyio_1.64.0
[13] DBI_1.1.2             lazyeval_0.2.2        BiocGenerics_0.40.0   colorspace_2.0-2
[17] withr_2.4.3           tidyselect_1.1.1      compiler_4.1.2        preprocessCore_1.56.0
[21] rvest_1.0.2           cli_3.2.0             Biobase_2.54.0        formatR_1.11
[25] xml2_1.3.3            plotly_4.10.0         scales_1.1.1          affy_1.72.0
[29] pbapply_1.5-0         digest_0.6.29         pkgconfig_2.0.3       htmltools_0.5.2
[33] dbplyr_2.1.1          fastmap_1.1.0         limma_3.50.0          htmlwidgets_1.5.4
[37] rlang_1.0.1           readxl_1.3.1          rstudioapi_0.13       impute_1.68.0
[41] generics_0.1.2        jsonlite_1.7.3        mzID_1.32.0           BiocParallel_1.28.3
[45] MALDIquant_1.21       Rcpp_1.0.8            munsell_0.5.0         S4Vectors_0.32.3
[49] fansi_1.0.2           MsCoreUtils_1.6.0     lifecycle_1.0.1       vsn_3.62.0
[53] stringi_1.7.6         MASS_7.3-55           zlibbioc_1.40.0       parallel_4.1.2
[57] crayon_1.5.0          lattice_0.20-45       haven_2.4.3           hms_1.1.1
[61] mzR_2.28.0            pillar_1.7.0          codetools_0.2-18      stats4_4.1.2
[65] futile.options_1.0.1  reprex_2.0.1          XML_3.99-0.8          glue_1.6.1
[69] lambda.r_1.2.4        pcaMethods_1.86.0     data.table_1.14.2     BiocManager_1.30.16
[73] modelr_0.1.8          vctrs_0.3.8           tzdb_0.2.0            foreach_1.5.2
[77] cellranger_1.1.0      gtable_0.3.0          clue_0.3-60           assertthat_0.2.1
[81] broom_0.7.12          ncdf4_1.19            viridisLite_0.4.0     iterators_1.0.14
[85] IRanges_2.28.0        cluster_2.1.2         ellipsis_0.3.2
```

**Reference**


1. Liang, L. *et al.* Metabolic Dynamics and Prediction of Gestational Age and Time to Delivery in Pregnant Women. *Cell* 181, 1680–1692.e15 (2020).
